# From manual counting to YOLO: Using computer vision to automate large-scale fecundity assays in *C. elegans*

**DOI:** 10.1101/2025.08.22.671778

**Authors:** Linyao Peng, Hongyi Shui, Anne Janisch, Sarah Hesse, Victoria Feist, Amanda Gibson

## Abstract

Fecundity measurements play a crucial role in life history research, providing insights into reproductive fitness, population dynamics, and environmental responses. In the model nematode *Caenorhabditis elegans*, fecundity assays are widely used to study development, aging, and genetic or environmental influences on reproduction. *C. elegans* hermaphrodites have large numbers of offspring (>100), so manual counting of viable offspring is time-consuming and susceptible to human error. Automated counting methods have the potential to enhance throughput, accuracy, and precision in data collection.

We applied computer vision to 9,972 images of broods from individual *C. elegans* hermaphrodites from several strains under multiple treatments to capture variation in fecundity. We trained models using YOLO versions v8 to v11 (large and extra-large variants) to detect and count viable offspring, then compared the model results to estimates from manual counting.

The best model was trained by YOLO v11-L. After fine-tuning, this model correctly detected 92% of all offspring visible in the images (recall) and was correct about 94.6% of the offspring it marked (precision). Manual counts differed from verified ground-truth counts by an average of 2.65 offspring per image, compared to 0.95 for the trained computer vision model. In addition, we detected significant effects of counter identity, experimental block, and their interaction on manual counts. Computer vision counts were not affected by these biases and outperformed manual counting in both speed and consistency.

We demonstrate that computer vision can be a powerful tool for fecundity assays in *C. elegans* and provide a pipeline for applying this approach to new image sets. More broadly, applying computer vision to digital collections can advance ecological and evolutionary research by accelerating the study of fitness and life history.

## Introduction

Fecundity assays are a fundamental approach in ecology and evolution (Charlesworth et al., 2007; Flatt, 2020; Flatt & Heyland, 2011; Knight & Robertson, 1957; Stearns, 1976). They give us insight into reproductive output, population dynamics, and fitness (Cheever et al., 1999; Miller, 2019; Pincheira-Donoso & Hunt, 2017; Witthames, PR et al., 2009). They are routinely used to measure the sensitivity of reproductive success to environmental stressors, including variation in temperature, pollution, parasitism, and exposure to chemicals (Charlesworth, 2015; González-Tokman et al., 2020; Mukai, 1964; Stearns, 2000).

Fecundity assays are especially popular in research with the model nematode *Caenorhabditis elegans*. An individual’s total reproductive output can be tracked over the course of ∼5 days and used to estimate key life history parameters (Boyd et al., 2010; Diaz & Viney, 2014; Galimov & Gems, 2020; Zhang et al., 2021). These studies tackle important questions on the impact of standing genetic variation, mutations, diet and environmental stressors promote or inhibit reproductive success (Davies & Hart, 2008; Prasad et al., 2011). *C. elegans* also serves as a model for investigating other life history traits that are tightly coupled to reproduction, such as developmental rate and lifespan (Johnson & Wood, 1982; Maklakov et al., 2017).

Traditionally, fecundity assays in *C. elegans* are conducted manually using dissecting microscopes to count the number of viable offspring deposited by an individual hermaphrodite.

*C. elegans* is highly fecund, producing 100-300 offspring quickly, within a few days (Brenner, 1974; Byerly et al., 1976). Thus, these assays provide rich data, but they are also labor-intensive and prone to human error (Hussey & Barker, 1973; Seinhorst, 1986; Waliullah et al., 2020). These features limit the use of fecundity assays at larger scales, especially in genome-wide association studies and multifactorial tests of environmental stressors. Large-scale fecundity assays have instead been carried out using high-throughput instruments such as the COPAS BIOSORT large particle sorter (Andersen et al., 2015) or WormScan (Mathew et al., 2012). Although these approaches enable powerful large-scale analyses, such platforms impose cost and accessibility barriers for many laboratories.

Computer vision may provide a cost-effective solution to *C. elegans* fecundity assays. Computer vision has already been used to quantify reproduction and morphology of various nematode species. Deep learning frameworks that utilized convolutional neural networks (CNNs) accurately identified and counted soybean cyst nematode (*Heterodera glycines*) eggs (Akintayo et al., 2018) and phenotyped beet cyst nematodes (*Heterodera schachtii*) (Chen et al., 2022). More recent efforts have applied YOLO (You Only Look Once)-based models to detect and classify eggs and juveniles of root knot nematode (*Meloidogyne*) species (Saikai et al., 2024) and to develop decision-support tools for estimating population sizes of plant-parasitic nematodes (Pun, Neupane, Koech, et al., 2023). These advancements underscore the potential for computer vision technologies to revolutionize nematode quantification, providing efficient and reliable alternatives to manual counting.

Computer vision models have been increasingly applied to automate counting in *C. elegans*. These models use a range of deep learning approaches, including convolutional neural networks (García Garví et al., 2021; White et al., 2013), YOLO versions (Akpu et al., 2025; Banerjee et al., 2023; Rico-Guardiola et al., 2022), Mask R-CNN (Fudickar et al., 2021; Rico-Guardiola et al., 2022), and vision transformers (Deserno & Bozek, 2023). They show high accuracy in detecting, classifying, and tracking individuals across developmental stages and during behavioral assays. The success of these prior models lays the groundwork for the application of computer vision in fecundity assays. Fecundity assays present particular challenges for existing computer vision models, especially when assays are conducted at scale using budget-friendly equipment. Images of broods may not be consistently high quality, offspring are rarely the same size, and they often overlap one another. Offspring are also typically imaged as early as possible, when they are very small, to ensure they are counted before dying, leaving the plate, or reproducing. Thus, we sought to develop a method for robust and accurate nematode counting that is compatible with variable image quality, flexible experimental set-ups, and mixed stage broods, including early life stages.

To accomplish this goal, we used YOLO versions to train models to count offspring in a large collection of images of *C. elegans* broods. We found that the computer vision models provided fast and reliable estimates of fecundity with less bias than manual counting. Our method can easily be modified for use by other nematode researchers.

## Materials and Methods

This section describes the complete workflow for training and evaluating computer vision models for automatic offspring counting in *C. elegans* fecundity assays. We describe our low-cost imaging setup to generate images, and our approach for designing a custom processing pipeline to isolate regions of interest and standardize image format. We then detail the protocol for annotating images, partitioning the dataset, training models using YOLO model versions, and fine-tuning the top-performing model. Finally, we describe statistical analyses for assessing model accuracy. A schematic of the full pipeline is shown in Figure 1. High resolution example photos are shown in Figure S1-4.

**Figure 1:**
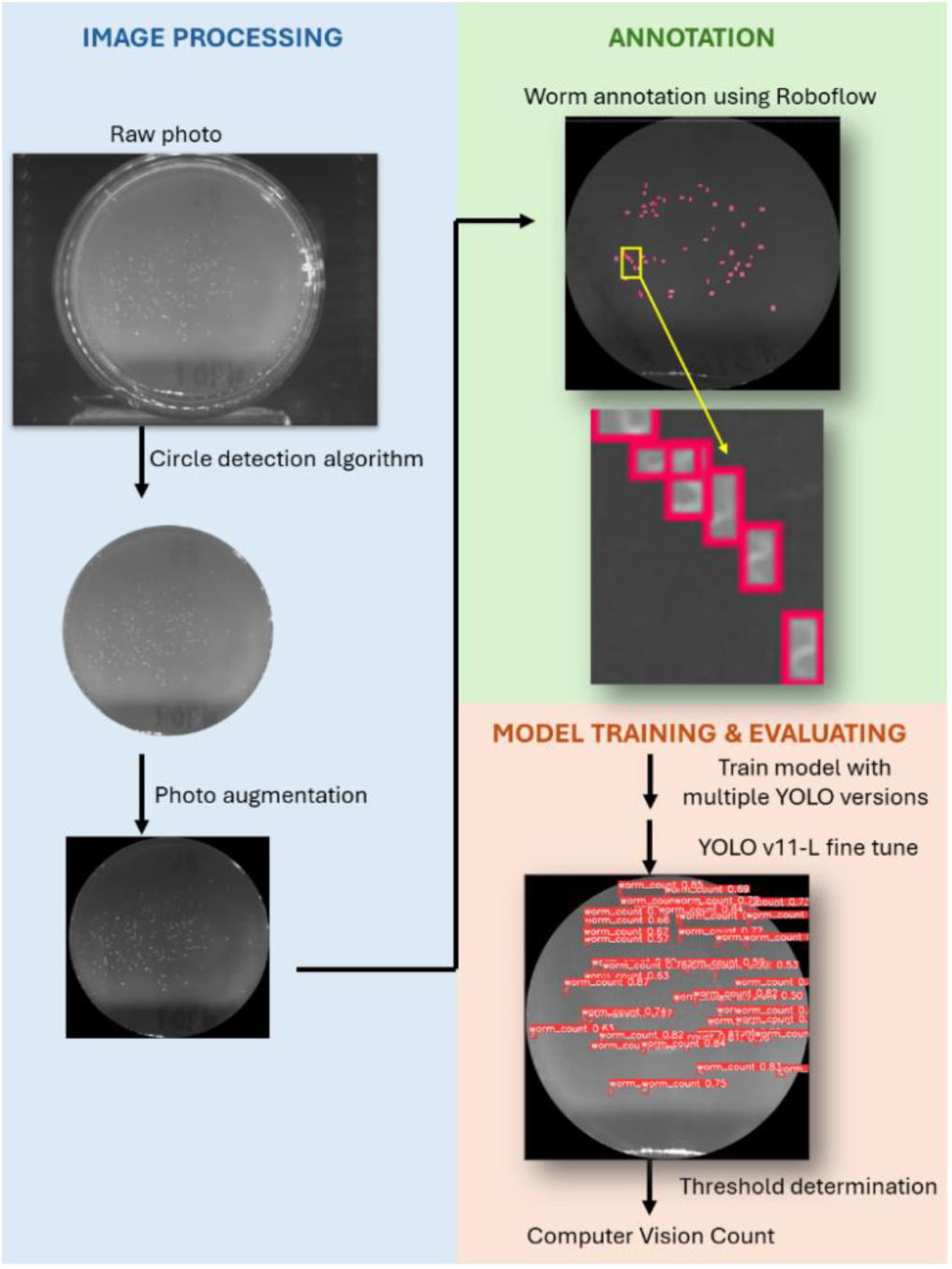
Overview of our workflow

### Image Acquisition and Dataset Preparation

#### Equipment

We built a low-cost imaging station adapted from Churgin and Fang-Yen (2015) and optimized for imaging fecundity plates based on Zhang et al. (2021). Imaging was performed using a DMK 33GP031monochrome GigE camera (The Imaging Source) with a 1/ 2.5’’ Aptina CMOS MT9P031 sensor (2592 X 1944 resolution). Two 1mm C/CS extension tubes (LAex5, LAex1) were used to adjust the optical path. Table S2 lists all the components, and Figure S5 shows the imaging station. This simple and affordable set up provided reliable and high-resolution images. This imaging station can be modified to provide higher-quality images, but we used this basic version to evaluate the potential for computer vision to perform with images that are low quality and less standardized.

#### Dataset Composition

High-resolution images of broods were captured under controlled laboratory conditions with standardized lighting, positioning, and imaging parameters to ensure consistency across samples. Each image was assigned a unique randomized identification code to facilitate tracking and subsequent annotation processes. The images were of groups of viable offspring hatched and reared on 35 mm petri dishes containing 4 mL of Nematode Growth Medium Lite (NGM Lite) seeded with 20 µL of *E. coli* OP50. Each petri dish was imaged twice to facilitate identification of offspring. Offspring were produced by individual hermaphrodites from different *C. elegans* strains that were either unexposed (control) or exposed to *Nematocida parisii* or *Nematocida ironsii*, each at high and low spore doses. The complete dataset comprised 9,972 high-resolution images acquired and stored in TIFF format. All images were hand-counted to generate manual count data for subsequent analyses.

#### Image Format and Pre-processing

We implemented a custom Python-based preprocessing pipeline to isolate the region of interest (ROI) containing the specimens through the following sequential steps:

1. **Circular Region Detection:** The Hough Circle Transform algorithm was employed to detect and isolate the circular petri dish in each image, establishing the region for subsequent analysis.
2. **Grayscale Conversion and Smoothing:** Images were converted to grayscale and smoothed using a 3 × 3 Gaussian blur kernel, which enhanced specimen-background contrast while reducing noise that could interfere with detection.
3. **ROI Extraction:** A binary mask was generated to isolate the petri dish region, eliminating background elements that could potentially introduce false positives during model inference.
4. **Normalization and Standardization:** The cropped images were resized to maintain dimensional consistency across all samples and saved in PNG format to preserve image quality while reducing file size.

This pre-processing pipeline ensured that YOLO models focused on specimen-containing regions while eliminating potential confounding background artifacts that could compromise detection accuracy.

### Image Annotation Protocol

A total of 331 images were randomly selected from the complete dataset and manually annotated to create a “ground-truth” image set for model training. For annotation, we used (*Roboflow*, n.d.) to label all offspring in images. These ground-truth labels were manually created bounding boxes that designate ‘true’ nematode. Bounding boxes were carefully delineated around each specimen, with particular attention to overlapping specimens, which were individually labeled to ensure comprehensive annotation of all visible organisms. To ensure consistency, a single annotator applied all labels. To maintain full control over the data preparation process, we only used Roboflow for annotation. We performed all preprocessing and augmentation of images independently in Python, as described above.

We exported the annotated dataset in PNG, a YOLO-compatible format, including both the image files and label files with normalized bounding box coordinates (center x, center y, width, height). We divided the ground-truth dataset into subsets to be used for model training (231 images, 70%), testing and hyperparameter optimization (50 images, 15%), and final model evaluation and performance assessment (50 images, 15%). To ensure balanced and unbiased subsets, we divided the dataset using randomized stratification, a process that preserves the distribution of specimen counts across subsets.

### Model Training and Fine-tuning

#### Initial YOLO Model Training

We systematically evaluated multiple pre-trained YOLO deep learning architectures for nematode detection and quantification (Redmon et al., 2016). We selected YOLO frameworks for this purpose because they can perform object localization and classification in a single forward pass of the neural network, making them particularly efficient for detection tasks in high-throughput biological imaging applications. The implementation was based on the Ultralytics framework with PyTorch (Paszke et al., 2019) as the underlying deep learning library.

We trained eight state-of-the-art large and extra-large YOLO model variants (YOLOv8-L/X, YOLOv9-C/E, YOLOv10-L/X, YOLOv11-L/X) under identical conditions to enable direct performance comparison. These models are part of the YOLO family of single-stage object detectors (Redmon et al., 2016), implemented through the Ultralytics framework (Jocher et al., 2023). Sapkota et al. (2025) provides a comprehensive review of YOLO developments through v11. All models were trained for 50 epochs with the following standardized hyperparameters: input image size of 1344 × 1344 pixels, batch size of 1, and 8 workers for data loading. The standard YOLO loss function, which incorporates localization, classification, and objectness components, was employed alongside the Adam optimizer with default YOLO hyperparameters. Training and evaluation of the YOLO models were conducted on a local workstation equipped with an NVIDIA GeForce RTX 2080 GPU (8GB VRAM) and 15.5 GB RAM. The software environment comprised Python 3.10.12, PyTorch 2.3.1, OpenCV, and CUDA 12.1. To enhance computational efficiency and training stability, batch normalization was applied to stabilize gradients, and mixed precision training was enabled to accelerate computation while maintaining numerical precision.

#### Confidence Threshold Optimization Using Five-Fold Cross-Validation

A critical parameter in object detection systems is the confidence threshold that determines which predictions are accepted as valid detections. For our purposes, this is the threshold at which an object is counted as a nematode. To systematically identify the optimal confidence threshold for each model, we implemented a rigorous five-fold cross-validation approach. The 50 images in the validation set were divided into five non-overlapping subsets, ensuring each fold contained images with a representative distribution of specimen counts to minimize bias in threshold selection. Confidence thresholds ranging from 0.01 to 1.0 were evaluated at fine-grained 0.01 increments. For each threshold value, the model-generated nematode count (cv_count) was compared against the ground truth count (gt_count). We used mean absolute error (MAE) as the primary optimization metric. MAE is calculated as the average absolute difference between predicted and ground truth counts for an image. For each fold in the five-fold cross-validation approach, we identified the confidence threshold that resulted in the minimum MAE between predicted and ground-truth counts (Figure 2). We then averaged these optimal thresholds across all five folds to determine a final threshold for each model. This approach helped lower the risk of overfitting to any single subset of the data and ensured that the chosen final threshold worked well across a range of image conditions (e.g., varying offspring density per plate).

**Figure 2.**
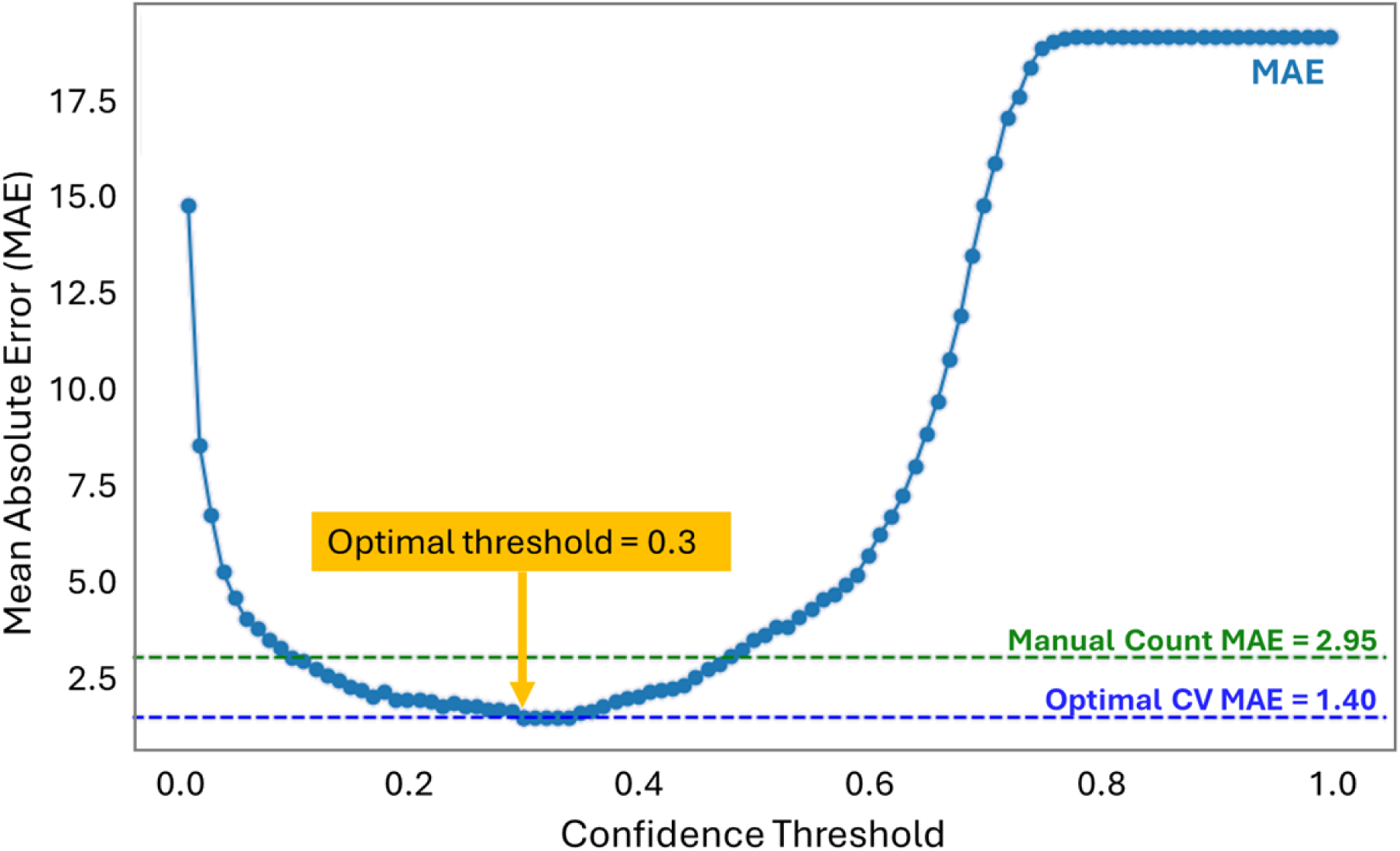
Example chart illustrating the confidence threshold selection process (shown for fold number 5). The green dashed line represents the mean absolute error (MAE) between manual counts and ground truth. The blue dashed line represents the optimal MAE between computer vision (CV) counts and ground truth. The optimal confidence threshold (yellow arrow) corresponds to the x-value at which MAE is minimized (blue dashed line), here at 0.3.

### Model Evaluation and Performance Metrics

Five-fold confidence threshold optimization allowed us to identify the optimal confidence threshold for each of the eight models trained by the different YOLO variants. We then assessed the detection capabilities of each model at its optimal confidence threshold using a variety of performance metrics to compare computer vision counts to ground annotations. We followed YOLO standard practice in selecting metrics that are relevant to our primary goal of accurate quantification (Redmon et al. 2016). The performance metrics included:

- **Precision (P)**: The proportion of correctly detected nematodes among all detections, calculated as: P = TP/(TP+FP) where TP represents true positives and FP represents false positives. This metric quantifies the model’s ability to avoid false detections.
- **Recall (R)**: The proportion of actual nematodes correctly detected, calculated as: R = TP/(TP+FN) where FN represents false negatives. This metric quantifies the model’s ability to detect all specimens present in the image.
- **F1-score**: The harmonic mean of precision (P) and recall (R), calculated as: F1 = 2×(P×R)/(P+R). This metric summarizes the trade-off between precision and recall, reflecting both the proportion of nematodes correctly detected and the extent to which true nematodes were missed or falsely identified.
- **Mean Absolute Error (MAE)**: The average absolute difference between predicted and ground truth counts for each image. This metric directly quantifies counting accuracy.

We used these metrics to identify the most suitable model for subsequent optimization.

### YOLOv11-L Model Fine-tuning

Based on comparison of these performance metrics, the model trained with the YOLOv11-L variant was selected for further optimization to enhance its overall detection capabilities. This step is referred to as “hyperparameterization.”

We selected values for key hyperparameters to fine-tune the YOLO model’s detection performance. These hyperparameters control the training process and were optimized to achieve the best balance between accuracy and computational efficiency. Table S1 presents the hyperparameters and their respective values evaluated during the fine-tuning process, along with the rationale for their selection.

The fine-tuning procedure was conducted on the YOLOv11-L model pre-trained during the initial training phase. To optimize detection performance, we tested multiple experimental configurations, each corresponding to a different combination of hyperparameter values. For each configuration, the model was trained for 50 epochs with consistent settings: input image dimensions of 1344 × 1344 pixels, batch size of 1, and 8 workers for data loading. A five-fold cross-validation approach was employed for confidence threshold optimization throughout the fine-tuning process to ensure robustness of the results and minimize sensitivity to data partitioning.

### Statistical Analyses

A strength of our data set is that each computer vision count is paired with a manual count of the same petri dish. This set up allows us to assess the accuracy and consistency of computer vision-based counts versus manual counts. Manual counts were conducted by three trained human counters. The computer vision counts were sourced from the fine-tuned Yolo v11-L model applied to each image. Each petri dish was imaged twice, so both images were analyzed by the model, and we used the rounded-up average of the two counts as the final value for analysis. Statistical analyses were performed in R (version 4.4.3) (R Core Team).

To assess the agreement between manual counts and computer vision counts, we calculated the correlation between the two sets of values using a simple linear regression. We further compared agreement and evaluated bias using a Bland-Altman analysis. For each image, we plotted the difference between the manual and computer vision counts against the average of the two methods. We then calculated the mean difference (bias) and the 95% limits of agreement, defined as the mean difference ± 1.96 times the standard deviation of the differences. This plot allowed us to visualize both the typical discrepancy between methods and the range within which most count differences fall.

Computer vision counts may be more accurate than manual counts, but perhaps equally important is their potential to be more consistent than manual counts. Consistency may especially be an issue for large assays with many images, such that manual counts must be conducted by multiple people or by a single person over multiple sessions. To assess consistency in our manual counts, we determined if mean bias differed significantly between counters, and whether the bias among counters was inconsistent (i.e., changed across blocks). We performed a two-way ANOVA with bias as the response variable and block, human counter, and their interaction as the predictors.

## Results

We first evaluated the performance of multiple YOLO variants to identify a suitable base model. We selected YOLOv11-L as the optimal variant and fine-tuned hyperparameters to improve performance. Finally, we compared computer vision counts to manual counts, revealing counter-specific and block-specific biases in manual counts.

### Initial YOLO Model Evaluation

We compared YOLO variants using precision, recall, F1-scores, and mean absolute error (MAE). The performance metrics for all evaluated YOLO variants are presented in Table 1. An example of standard output performance results from the YOLO model is shown in Figure S6. Among the eight models initially evaluated, YOLOv11-L demonstrated consistently strong performance across metrics, ranking in the top three across all metrics. Notably both precision (0.924) and recall (0.890) were high, resulting in a strong F1-score (0.907) and low MAE (1.17). For our purposes, both over- and under-counting are problematic, so it is critical to achieve a balance between precision, which minimizes false positives, and recall, which minimizes false negatives.

**Table 1:**
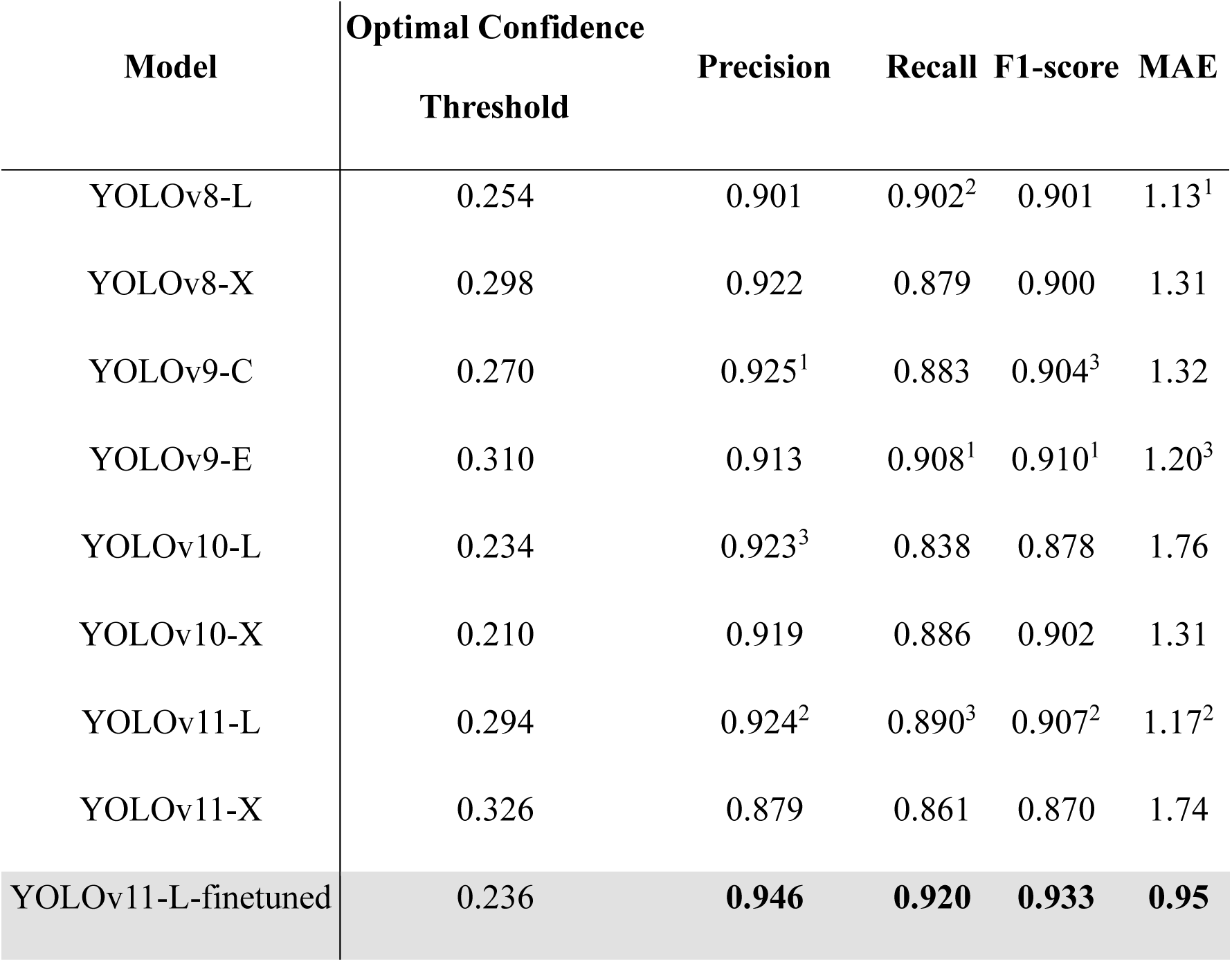
Performance metrics for trained models for each YOLO variant, as well as the fine-tuned YOLOv11-L model. Subscripts indicate the three top scores for each performance metric among the initial models. The fine-tuned model outperforms all initial models.

Other models performed well in some metrics, but not all. For example, YOLOv8-L achieved high recall (0.902) and the best MAE (1.13), but its precision (0.901) was substantially lower than YOLOv11-L. Similarly, YOLOv9-E achieved better recall (0.908) and a slightly better F1-score (0.910), but its precision (0.913) and MAE (1.20) were substantially lower.

We found that the larger variants of YOLO ((i.e., the high-capacity “X” models with more layers and parameters) did not consistently outperform their lighter (“L”) counterparts despite their increased capacity. For instance, YOLOv11-X performed worse than YOLOv11-L across all metrics. This result aligns with previous findings: newer or larger YOLO models do not always surpass older versions in real-world tasks (Jiang & Zhong, 2025), and optimizations often yield larger gains than scaling model size alone (Ali, Bhowmik & Nicol,.2023).

### Fine-tuning

After identifying YOLOv11-L as the most promising variant, we conducted systematic hyperparameter optimization to further enhance its detection capabilities. We tested 192 distinct hyperparameter combinations (2 optimizers × 4 IoU thresholds × 4 HSV_V values × 4 scale factors × 4 perspective coefficients × 4 mosaic probabilities) using the same five-fold cross-validation protocol employed during the initial model assessment (Table S1).

We identified the optimal configuration to be: SGD (stochastic gradient descent) optimizer with IoU=0.4, HSV_V=0.5, scale=0.2, perspective=0.001, and mosaic=0.3. This fine-tuned YOLOv11-L model demonstrated substantial improvements across all metrics (Table 1). Precision, recall, and F1-score each increased by 2–3%, indicating fewer false positives and fewer missed nematodes. MAE also decreased substantially, from 1.17 to 0.95, reflecting an increase in overall counting accuracy. The optimized confidence threshold for the fine-tuned model (0.236) was lower than that of the base YOLOv11-L model (0.294), suggesting that the improved feature representation enabled more reliable detections at lower confidence scores.

This threshold reduction, combined with improved precision and recall, suggests that the fine-tuned model more effectively distinguishes true nematodes from background elements.

### Comparison of computer vision and manual counts

In the validation set of annotated images (N=100), manual counts and computer vision counts showed similar deviations from ground truth counts (Figure 3). The mean absolute error (MAE) for manual counts was 2.65 nematodes per image, compared to 0.95 nematodes per image for the fine-tuned YOLOv11-L predictions. While manual counts exhibited a wider range of both over- and undercounting, the overall average error did not differ significantly between the two methods in this dataset (two-sided t-test: test-stat; df= 0, p=0.14).

**Figure 3.**
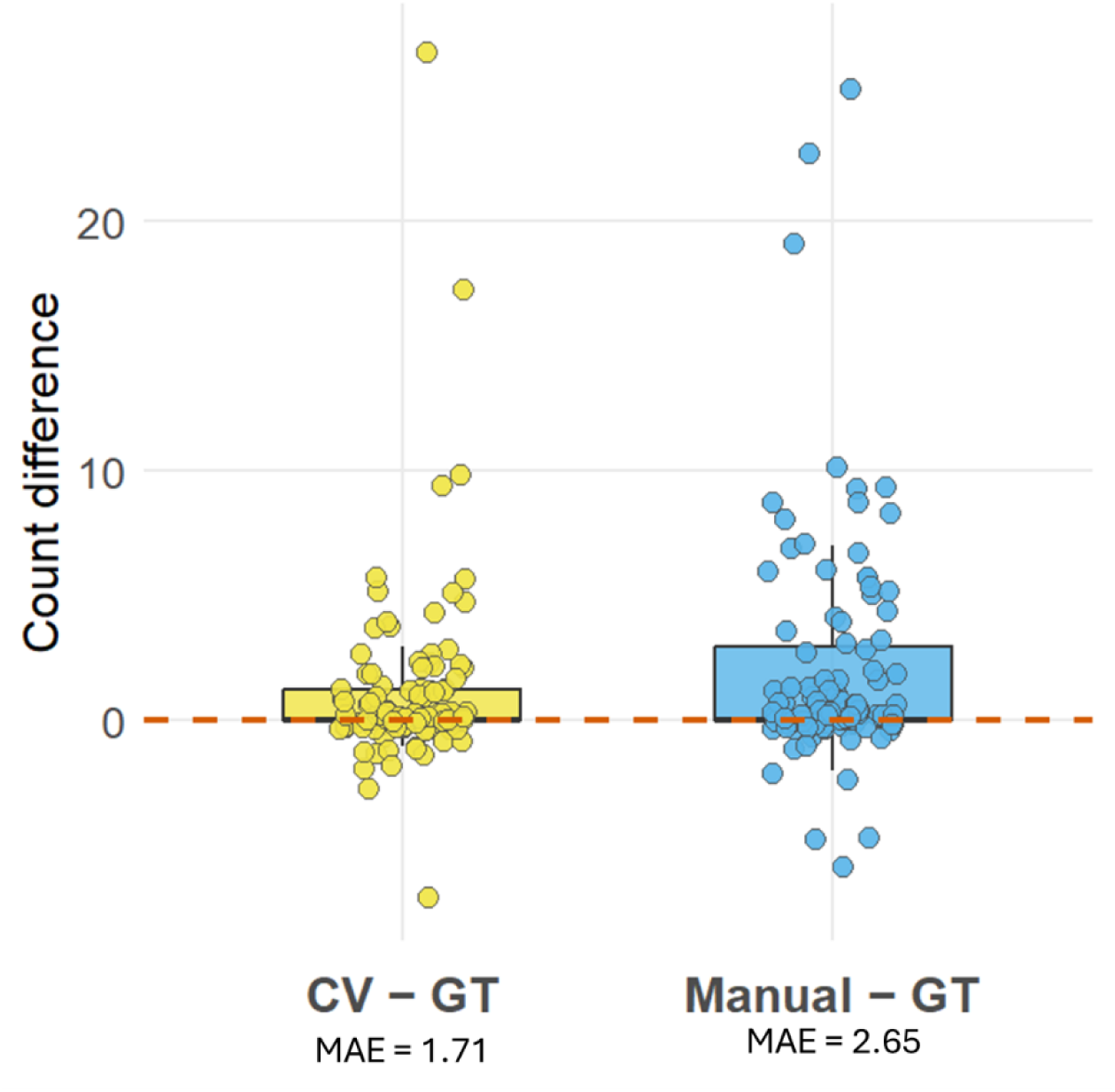
Differences from ground truth counts (GT) for computer vision (CV) and manual counts in the evualuation set of annotated images. Computer vision counts reflect the fine-tuned YOLOv11-L model predictions. Each point represents one image; boxes show interquartile ranges. The dashed line at zero indicates perfect agreement with the ground truth count of an image.

Consistent with this finding, the fine-tuned YOLOv11-L counts showed a strong linear relationship with manual counts (linear regression: R² = 0.93; Figure 4A), indicating high consistency between the two methods across the range of observed values. Bland-Altman analysis revealed a mean difference of 0.32 nematodes between manual and computer vision counts, suggesting minimal systematic bias (Figure 4B). The 95% limits of agreement ranged from −15.36 to 16.00, indicating variability in the differences between methods. Notably, at higher count values, many points fell outside the agreement bounds, although these cases represented a relatively small proportion of the dataset. We examined the 84 images with count discrepancies greater than 21 between manual and computer vision estimates. In 61 cases, the CV counts were more accurate than the manual counts. The remaining 23 images were compromised by either mixed generations of nematodes or poor photo quality.

**Figure 4.**
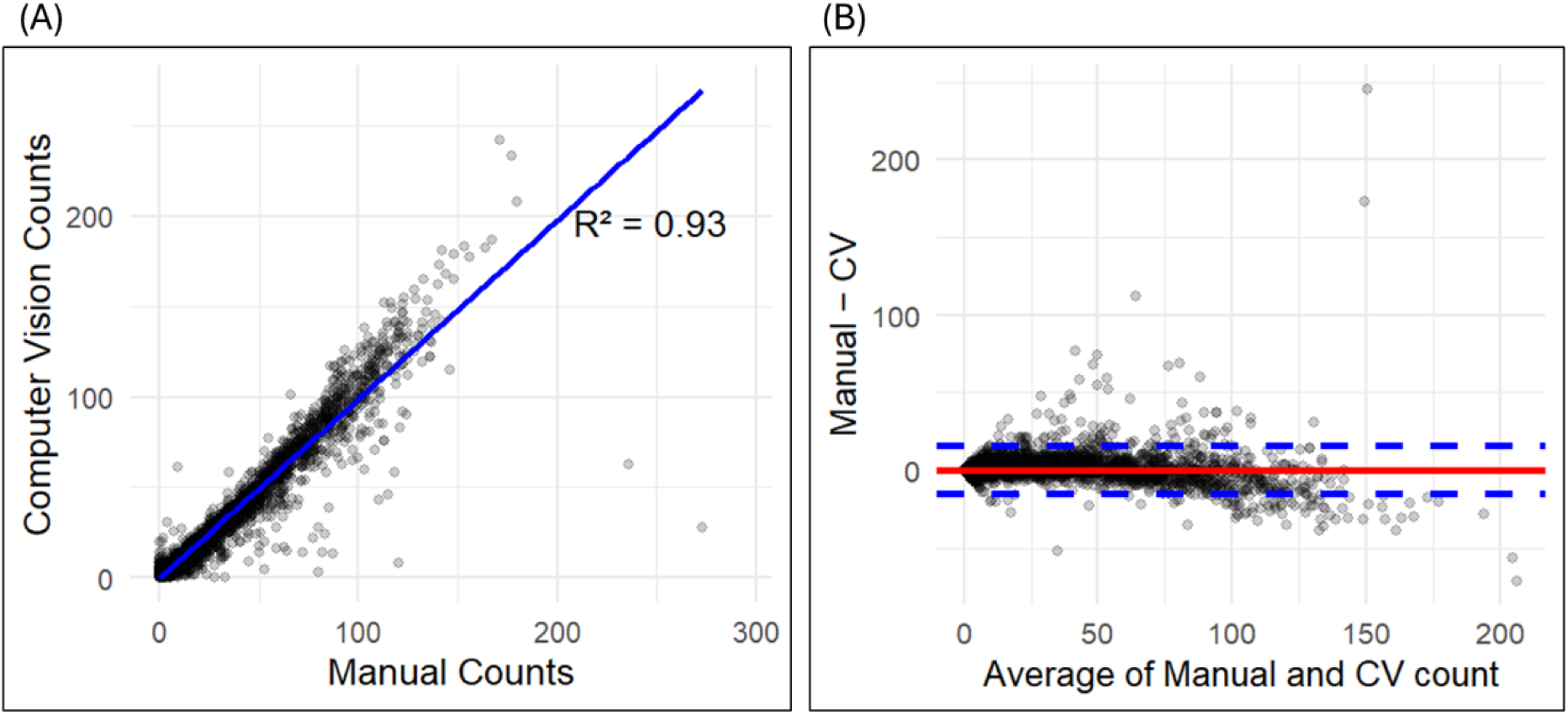
Comparison of manual and computer vision counts. (A) Correlation of computer vision counts and manual counts. Each point represents a single image, and the blue line shows the best-fit linear regression. (B) Bland-Altman plot showing agreement between manual and computer vision counts. Each point represents the difference between a manual count and a computer vision count, plotted against their average. The solid red line indicates the mean difference, and the dashed blue lines represent the 95% limits of agreement.

We observed significant effects of counter identity, experimental block, and their interaction on the bias between manual and computer vision counts (Figure 5). A two-way ANOVA revealed main effects of Counter (F(2, 5013) = 14.83, p = 3.78 × 10⁻⁷) and Block (F(5, 5013) = 15.82, p = 1.72 × 10⁻¹⁵), as well as a highly significant Counter × Block interaction (F(10, 5013) = 11.83, p = 2.03 × 10⁻²⁰). Post hoc comparisons showed that Counter #1 consistently overcounted by an average of 2.09 nematodes per image (95% CI: [1.62, 2.56], p < 0.001), whereas Counters #2 and #3 undercounted by −0.37 and −0.67 nematodes per image, respectively (p < 0.05 for both) (Figure 5A). The interaction effect indicates that these patterns of counter bias varied across blocks, which were counted in different weeks over a six-month period (Figure 5B). Together, these results demonstrate that YOLOv11-L-derived counts closely matched to manual counts while offering improved consistency.

**Figure 5.**
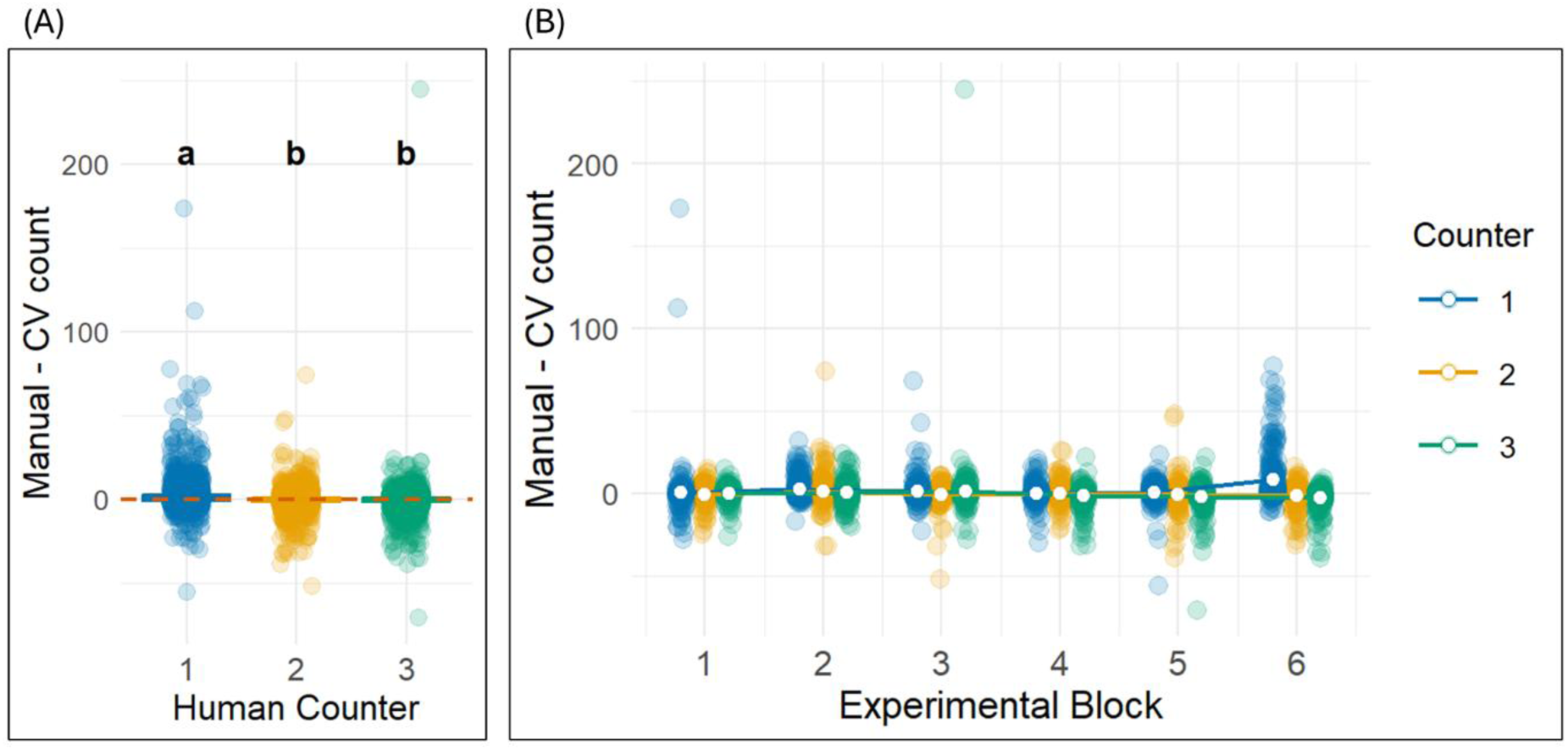
Bias between computer vision and manual nematode counts as a function of counter and block. (A) Counter bias: The difference between manual and computer vision (CV) counts for each human counter, with letters indicating Tukey HSD post hoc comparisons among counters. Each point represents a single image, and the dashed red line at zero denotes perfect agreement between methods. (B) Bias across experimental blocks, showing individual data points and mean trends for each counter.

## Discussion

In this study, we developed a computer vision pipeline based on the YOLOv11-L deep learning model to count viable offspring in *Caenorhabditis elegans* fecundity assays. Our approach represents a significant innovation by applying object detection to whole-plate images captured using a cheap and user-friendly camera setup. Our model was evaluated against a large, manually curated dataset, enabling a more rigorous comparison to traditional human assessment. We found that our computer vision model performed substantially better than manual counting.

### Advancing life history study in *C. elegans* using computer vision

The most evident advantage of our computer vision model is speed. With the development of machine learning models, computer vision–based object counting has become a powerful approach that dramatically accelerates large-scale assays (Babu et al., 2023; Kataras et al., 2023; McGuire et al., 2021; Vieira et al., 2025; Xu et al., 2023). In our project, manual counting of offspring was conducted incrementally over approximately six months, with each experimental block annotated in separate weeks. In contrast, computer vision-based counting was completed within a few days, including preprocessing, model inference, and post-processing. This dramatic difference highlights the value of automated approaches for high-throughput quantification. The framework is simple to train and use, and we provide full documentation and guidelines for adapting the approach to other systems here: https://github.com/Coevolving-Lab/nematode-fecundity-computer-vision-counting.

We also found that the YOLOv11-L model performs well. It achieved high precision and recall across a wide range of photos. Moreover, computer vision approaches are consistent within and between projects. In our analysis, computer vision counts avoided the variation that arose among multiple human counters. Large datasets typically require multiple counters, and variation among counters can be especially hard to account for if individual counters subtly modify their approach over time. Our computer vision approach thus has the potential to increase throughput and standardize data collection, improving the efficiency and quality of *C. elegans* fecundity measurements. Beyond our immediate experimental system, the architecture and methodology are generalizable, offering potential for automated quantification of other microscopic organisms or phenotypic traits in biological imaging datasets.

### Limitations in Current *C. elegans* Life History Research

Early computational efforts with *C. elegans* largely focused on resolving clusters of overlapping nematodes using model-search algorithms (Raviv et al., 2010; Wählby et al., 2010, 2012). With advances in deep learning, particularly convolutional neural networks (CNNs), more recent pipelines have begun incorporating these architectures into nematode detection tasks.

Among these, the YOLO family of models has gained prominence for its speed and accuracy in object detection since its introduction in 2015 (Matsuzaka & Yashiro, 2023; Zhao et al., 2019). Thus far, applications of YOLO in *C. elegans* research have primarily focused on behavioral and locomotion analyses (Banerjee et al., 2023; B. Dong & Chen, 2025; Mori et al., 2022), rather than quantification in static images. In contrast, object detection models such as YOLO have been widely used in studies of plant-parasitic nematodes, particularly root-knot and cyst nematodes. These models enable automated classification and quantification of nematodes for use in field diagnostics and agricultural management (Pun, Neupane, & Koech, 2023). This difference in the application of computer vision among nematode models likely reflects research priorities: in agricultural systems, numbers and life history traits of pests directly impact crop yield, driving demand for high-throughput quantification tools.

Broader application of computer vision in life history assays would advance *C. elegans* as a system for addressing key ecological and evolutionary questions. Our dataset pushed the model to perform under realistic and variable biological settings, thereby supporting its application in life history experiments. Our images varied in offspring number, object clustering, and background noise. Offspring counts ranged from dozens to hundreds, with overlapping bodies and subtle morphological variation. Our success given this wide range of image conditions highlights the flexibility of YOLOv11-L to operate in diverse and noisy environments. As such, we hope our pipeline not only bridges a methodological gap but also catalyzes broader adoption of computer vision in *C. elegans* reproductive biology.

### Challenges and Future Opportunities

While our model showed strong performance, several challenges remain. First, the resolution and quality of images imposed upper limits on detection accuracy. Some photos in our dataset were downsampled due to file size constraints, reducing the model’s ability to resolve tightly clustered or low-contrast individuals. Similar effects of image resolution on object detection performance have been reported in other contexts (Hao et al., 2023). Future work could address this by integrating higher-resolution cameras, refining imaging pipelines, or applying super-resolution techniques during preprocessing. These approaches has been shown to enhance small object detection in computer vision tasks (Ardohain & Fei, 2025; C. Dong et al., 2015; Yang et al., 2019). However, we used this low-cost imaging setup to reduce barriers to large-scale fecundity assays, and our model achieved high accuracy nonetheless, demonstrating the practical viability of this approach.

Second, improving accessibility was one of our key goals. One distinct advantage of our approach is its ability to run on a standard desktop workstation, avoiding the need for high-end computational resources. This contrasts with many large-scale object detection applications, which typically rely on commercial servers or cloud platforms to handle training and inference at scale (Shankar et al., 2020; Tuli et al., 2019; Yaseen et al., 2018). By eliminating this barrier, our method is more practical for widespread use in academic laboratories with limited infrastructure. To further enhance accessibility, we envision packaging the model into a user-friendly graphical user interface (GUI), enabling researchers without coding experience to upload images, perform automated analyses, and export fecundity estimates. Such a tool would significantly lower the barrier to application, particularly in life history and evolutionary biology studies using *C. elegans*.

## Conclusion

Together, our results demonstrate the feasibility and value of automated object detection in quantifying reproductive output in *C. elegans*. By combining deep learning with high-resolution life history data, we present a scalable and accurate alternative to manual counting.

This work lays the groundwork for future integration of computer vision into fecundity assays and opens the door to more standardized and comprehensive analyses of life history traits across experimental systems.

## Acknowledgements

This study was funded by the US National Institutes of Health MIRA Award to Amanda K. Gibson (5R35GM137975). The authors thank Erik Andersen (Johns Hopkins University, Department of Biology) for assistance with the camera setup, and all members of the Gibson Lab for their helpful feedback. We are also grateful to Keric Lamb (University of Virginia, Department of Biology) for recommending the statistical analysis used to compare the two counting methods.

## Data Availability Statement

Components of the camera imaging station setup are in Table S2. The data that supports the findings of this study are available within the article and its Supplementary files. The entire pipeline of the computer vision model is available at the GitHub repository https://github.com/Coevolving-Lab/nematode-fecundity-computer-vision-counting.

## References

Akintayo, A., Tylka, G. L., Singh, A. K., Ganapathysubramanian, B., Singh, A., & Sarkar, S. (2018). A deep learning framework to discern and count microscopic nematode eggs. Scientific Reports, 8(1), 9145. 10.1038/s41598-018-27272-w

Akpu, C. H., Wei, H., & Hong, X. (2025). Deep Learning in Automated Worm Identification and Tracking for *C. Elegan* Mating Behavior Analysis. In: Antonacopoulos, A., Chaudhuri, S., Chellappa, R., Liu, CL., Bhattacharya, S., Pal, U. (eds) Pattern Recognition. ICPR 2024. Lecture Notes in Computer Science, vol 15303. Springer, Cham. 10.1007/978-3-031-78122-3_8

Andersen, E. C., Shimko, T. C., Crissman, J. R., Ghosh, R., Bloom, J. S., Seidel, H. S., Gerke, J. P., & Kruglyak, L. (2015). A powerful new quantitative genetics platform, combining *Caenorhabditis elegans* high-throughput fitness assays with a large collection of recombinant strains. G3 Genes|Genomes|Genetics, 5(5), 911–920. 10.1534/g3.115.017178

Ardohain, C., & Fei, S. (2025). The impacts of training data spatial resolution on deep learning in remote sensing. Science of Remote Sensing, 11, 100185. 10.1016/j.srs.2024.100185

Babu, K. M., Bentall, D., Ashton, D. T., Puklowski, M., Fantham, W., Lin, H. T., Tuckey, N. P. L., Wellenreuther, M., & Jesson, L. K. (2023). Computer vision in aquaculture: A case study of juvenile fish counting. Journal of the Royal Society of New Zealand, 53(1), 52–68. 10.1080/03036758.2022.2101484

Banerjee, S. C., Khan, K. A., & Sharma, R. (2023). Deep-worm-tracker: Deep learning methods for accurate detection and tracking for behavioral studies in *C. elegans*. Applied Animal Behaviour Science, 266, 106024. 10.1016/j.applanim.2023.106024

Boyd, W. A., McBride, S. J., Rice, J. R., Snyder, D. W., & Freedman, J. H. (2010). A high-throughput method for assessing chemical toxicity using a *Caenorhabditis elegans* reproduction assay. Toxicology and Applied Pharmacology, 245(2), 153–159. 10.1016/j.taap.2010.02.014

Brenner, S. (1974). The genetics of *Caenorhabditis elegans*. Genetics, 77(1), 71–94. 10.1093/genetics/77.1.71

Byerly, L., Cassada, R. C., & Russell, R. L. (1976). The life cycle of the nematode *Caenorhabditis elegans*. Developmental Biology, 51(1), 23–33. 10.1016/0012-1606(76)90119-6

Charlesworth, B. (2015). Causes of natural variation in fitness: Evidence from studies of *Drosophila* populations. Proceedings of the National Academy of Sciences, 112(6), 1662– 1669. 10.1073/pnas.1423275112

Charlesworth, B., Miyo, T., & Borthwick, H. (2007). Selection responses of means and inbreeding depression for female fecundity in *Drosophila melanogaster* suggest contributions from intermediate-frequency alleles to quantitative trait variation. Genetical Research, 89(2), 85–91. 10.1017/S001667230700866X

Cheever, A. W., Poindexter, R. W., & Wynn, T. A. (1999). Egg laying is delayed but worm fecundity is normal in SCID mice infected with *Schistosoma japonicum* and *S. mansoni* with or without recombinant tumor necrosis factor alpha treatment. Infection and Immunity, 67(5), 2201–2208. 10.1128/IAI.67.5.2201-2208.1999

Chen, L., Daub, M., Luigs, H.-G., Jansen, M., Strauch, M., & Merhof, D. (2022). High-throughput phenotyping of nematode cysts. Frontiers in Plant Science, 13, 965254. 10.3389/fpls.2022.965254

Churgin, M. A., & Fang-Yen, C. (2015). An imaging system for *C. elegans* behavior. In D. Biron & G. Haspel (Eds.), C. elegans (Vol. 1327, pp. 199–207). Humana Press. 10.1007/978-1-4939-2842-2_14

Davies, K. G., & Hart, J. E. (2008). Fecundity and lifespan manipulations in *Caenorhabditis elegans* using exogenous peptides. Nematology, 10(1), 103–112. 10.1163/156854108783360221

Deserno, M., & Bozek, K. (2023). WormSwin: Instance segmentation of *C. elegans* using vision transformer. Scientific Reports, 13(1), 11021. 10.1038/s41598-023-38213-7

Diaz, S. A., & Viney, M. (2014). Genotypic-specific variance in *Caenorhabditis elegans* lifetime fecundity. Ecology and Evolution, 4(11), 2058–2069. 10.1002/ece3.1057

Dong, B., & Chen, W. (2025). A high precision method of segmenting complex postures in *Caenorhabditis elegans* and deep phenotyping to analyze lifespan. Scientific Reports, 15(1), 8870. 10.1038/s41598-025-93533-0

Dong, C., Loy, C. C., He, K., & Tang, X. (2015). Image super-resolution using deep convolutional networks (No. arXiv:1501.00092). *arXiv*. 10.48550/arXiv.1501.00092

Flatt, T. (2020). Life-history evolution and the genetics of fitness components in *Drosophila melanogaster*. Genetics, 214(1), 3–48. 10.1534/genetics.119.300160

Flatt, T., & Heyland, A. (Eds.). (2011). Mechanisms of life history evolution: The genetics and physiology of life history traits and trade-offs. Oxford University Press. 10.1093/acprof:oso/9780199568765.001.0001

Fudickar, S., Nustede, E. J., Dreyer, E., & Bornhorst, J. (2021). Mask R-CNN based *C. elegans* detection with a DIY microscope. Biosensors, 11(8), 257. 10.3390/bios11080257

Galimov, E. R., & Gems, D. (2020). Shorter life and reduced fecundity can increase colony fitness in virtual *Caenorhabditis elegans*. Aging Cell, 19(5), e13141. 10.1111/acel.13141

García Garví, A., Puchalt, J. C., Layana Castro, P. E., Navarro Moya, F., & Sánchez-Salmerón, A.-J. (2021). Towards lifespan automation for *Caenorhabditis elegans* based on deep learning: Analysing convolutional and recurrent neural networks for dead or live classification. Sensors, 21(14), 4943. 10.3390/s21144943

González-Tokman, D., Córdoba-Aguilar, A., Dáttilo, W., Lira-Noriega, A., Sánchez-Guillén, R. A., & Villalobos, F. (2020). Insect responses to heat: Physiological mechanisms, evolution and ecological implications in a warming world. Biological Reviews, 95(3), 802–821. 10.1111/brv.12588

Hao, Y., Pei, H., Lyu, Y., Yuan, Z., Rizzo, J.-R., Wang, Y., & Fang, Y. (2023). Understanding the impact of image quality and distance of objects to object detection performance. 2023 IEEE/RSJ International Conference on Intelligent Robots and Systems (IROS), 11436–11442. 10.1109/IROS55552.2023.10342139

Hussey, R. S., & Barker, K. R. (1973). A comparison of methods of collecting inocula of *Meloidogyne spp.*, including a new technique. Plant Disease Reporter, 57(12), 1025–1028.

Jocher, G., Qiu, J., & Chaurasia, A. (2023). Ultralytics YOLO (Version 8.0.0) [Computer software]. Ultralytics. https://github.com/ultralytics/ultralytics

Johnson, T. E., & Wood, W. B. (1982). Genetic analysis of life-span in *Caenorhabditis elegans*. Proceedings of the National Academy of Sciences, 79(21), 6603–6607. 10.1073/pnas.79.21.6603

Kataras, T. J., Jang, T. J., Koury, J., Singh, H., Fok, D., & Kaul, M. (2023). ACCT is a fast and accessible automatic cell counting tool using machine learning for 2D image segmentation. Scientific Reports, 13(1), 8213. 10.1038/s41598-023-34943-w

Knight, G. R., & Robertson, A. (1957). Fitness as a measurable character in Drosophila. Genetics, 42(4), 524–530. 10.1093/genetics/42.4.524

Maklakov, A. A., Carlsson, H., Denbaum, P., Lind, M. I., Mautz, B., Hinas, A., & Immler, S. (2017). Antagonistically pleiotropic allele increases lifespan and late-life reproduction at the cost of early-life reproduction and individual fitness. Proceedings of the Royal Society B: Biological Sciences, 284(1856), 20170376. 10.1098/rspb.2017.0376

Mathew, M. D., Mathew, N. D., & Ebert, P. R. (2012). WormScan: A technique for high-throughput phenotypic analysis of *Caenorhabditis elegans*. PLoS ONE, 7(3), e33483. 10.1371/journal.pone.0033483

Matsuzaka, Y., & Yashiro, R. (2023). AI-based computer vision techniques and expert systems. AI, 4(1), 289–302. 10.3390/ai4010013

McGuire, M., Soman, C., Diers, B., & Chowdhary, G. (2021). High throughput soybean pod-counting with in-field robotic data collection and machine-vision based data analysis (No. arXiv:2105.10568). *arXiv*. 10.48550/arXiv.2105.10568

Miller, Z. (2019). Digest: Does sexual conflict complicate a trade-off between fecundity and survival?. Evolution, 73(11), 2347–2348. 10.1111/evo.13855

Mori, S., Tachibana, Y., Suzuki, M., & Harada, Y. (2022). Automatic worm detection to solve overlapping problems using a convolutional neural network. Scientific Reports, 12(1), 8521. 10.1038/s41598-022-12576-9

Mukai, T. (1964). The Genetic Structure of Natural Populations of *Drosophila Melanogaster*. I. Spontaneous Mutation Rate of Polygenes Controlling Viability. Genetics, 50(1), 1–19. 10.1093/genetics/50.1.1

Paszke, A., Gross, S., Massa, F., Lerer, A., Bradbury, J., Chanan, G., Killeen, T., Lin, Z., Gimelshein, N., Antiga, L., Desmaison, A., Köpf, A., Yang, E., DeVito, Z., Raison, M., Tejani, A., Chilamkurthy, S., Steiner, B., Fang, L., … Chintala, S. (2019). PyTorch: An imperative style, high-performance deep learning library (No. arXiv:1912.01703). *arXiv.* 10.48550/arXiv.1912.01703

Pincheira-Donoso, D., & Hunt, J. (2017). Fecundity selection theory: Concepts and evidence: Fecundity selection. Biological Reviews, 92(1), 341–356. 10.1111/brv.12232

Prasad, A., Croydon-Sugarman, M. J. F., Murray, R. L., & Cutter, A. D. (2011). Temperature-dependent fecundity associates with latitude in *Caenorhabditis briggsae*. Evolution, 65(1), 52–63. 10.1111/j.1558-5646.2010.01110.x

Pun, T. B., Neupane, A., & Koech, R. (2023). A Deep Learning-based decision support tool for plant-parasitic nematode management. Journal of Imaging, 9(11), 240. 10.3390/jimaging9110240

Pun, T. B., Neupane, A., Koech, R., & Walsh, K. (2023). Detection and counting of root-knot nematodes using YOLO models with mosaic augmentation. Biosensors and Bioelectronics: X, 15, 100407. 10.1016/j.biosx.2023.100407

Raviv, T. R., Ljosa, V., Conery, A. L., Ausubel, F. M., Carpenter, A. E., Golland, P., & Wählby, C. (2010). Morphology-guided graph search for untangling objects: *C. elegans* analysis. In T. Jiang, N. Navab, J. P. W. Pluim, & M. A. Viergever (Eds.), Medical Image Computing and Computer-Assisted Intervention – MICCAI 2010 (pp. 634–641). Springer. 10.1007/978-3-642-15711-0_79

Redmon, J., Divvala, S., Girshick, R., & Farhadi, A. (2016). You Only Look Once: Unified, real-time object detection. 2016 IEEE Conference on Computer Vision and Pattern Recognition (CVPR), 779–788. 10.1109/CVPR.2016.91

Rico-Guardiola, E. J., Layana-Castro, P. E., García-Garví, A., & Sánchez-Salmerón, A.-J. (2022). *Caenorhabditis Elegans* detection using YOLOv5 and Faster R-CNN Networks. In A. I. Pereira, A. Košir, F. P. Fernandes, M. F. Pacheco, J. P. Teixeira, & R. P. Lopes (Eds.), Optimization, Learning Algorithms and Applications (Vol. 1754, pp. 776–787). Springer International Publishing. 10.1007/978-3-031-23236-7_53

Roboflow. (n.d.). Roboflow: Computer vision dataset management, preprocessing, and model deployment platform. https://roboflow.com

Saikai, K. K., Bresilla, T., Kool, J., De Ruijter, N. C. A., Van Schaik, C., & Teklu, M. G. (2024). Counting nematodes made easy: Leveraging AI-powered automation for enhanced efficiency and precision. Frontiers in Plant Science, 15, 1349209. 10.3389/fpls.2024.1349209

Sapkota, R., Flores-Calero, M., Qureshi, R., Badgujar, C., Nepal, U., Poulose, A., Zeno, P., Vaddevolu, U. B. P., Khan, S., Shoman, M., Yan, H., & Karkee, M. (2025). YOLO advances to its genesis: A decadal and comprehensive review of the You Only Look Once (YOLO) series. Artificial Intelligence Review, 58(9), 274. 10.1007/s10462-025-11253-3

Seinhorst, J. W. (1986). The development of individuals and populations of cyst nematodes on plants. In F. Lamberti & C. E. Taylor (Eds.), Cyst Nematodes (pp. 101–117). Springer US. 10.1007/978-1-4613-2251-1_5

Shankar, K., Wang, P., Xu, R., Mahgoub, A., & Chaterji, S. (2020). JANUS: Benchmarking commercial and open-source cloud and edge platforms for object and anomaly detection workloads (No. arXiv:2012.04880). arXiv. 10.48550/arXiv.2012.04880

Stearns, S. C. (1976). Life-history tactics: A review of the ideas. The Quarterly Review of Biology, 51(1), 3–47. 10.1086/409052

Stearns, S. C. (2000). Life history evolution: Successes, limitations, and prospects. Naturwissenschaften, 87(11), 476–486. 10.1007/s001140050763

Tuli, S., Basumatary, N., & Buyya, R. (2019). EdgeLens: Deep learning based object detection in integrated IoT, Fog and cloud computing environments (No. arXiv:1906.11056). arXiv. 10.48550/arXiv.1906.11056

Vieira, A. B., Valente, M., Montezuma, D., Albuquerque, T., Ribeiro, L., Oliveira, D., Monteiro, J., Gonçalves, S., Pinto, I. M., Cardoso, J. S., & Oliveira, A. L. (2025). CountPath: Automating fragment counting in digital pathology (No. arXiv:2503.10520). *arXiv*. 10.48550/arXiv.2503.10520

Wählby, C., Kamentsky, L., Liu, Z. H., Riklin-Raviv, T., Conery, A. L., O’Rourke, E. J., Sokolnicki, K. L., Visvikis, O., Ljosa, V., Irazoqui, J. E., Golland, P., Ruvkun, G., Ausubel, F. M., & Carpenter, A. E. (2012). An image analysis toolbox for high-throughput *C. elegans* assays. Nature Methods, 9(7), 714–716. 10.1038/nmeth.1984

Wählby, C., Riklin-Raviv, T., Ljosa, V., Conery, A. L., Golland, P., Ausubel, F. M., & Carpenter, A. E. (2010). Resolving clustered worms via probabilistic shape models. 2010 IEEE International Symposium on Biomedical Imaging: From Nano to Macro, 552–555. 10.1109/ISBI.2010.5490286

Waliullah, S., Bell, J., Jagdale, G., Stackhouse, T., Hajihassani, A., Brenneman, T., & Ali, Md. E. (2020). Rapid detection of pecan root-knot nematode, *Meloidogyne partityla*, in laboratory and field conditions using loop-mediated isothermal amplification. PLOS ONE, 15(6), e0228123. 10.1371/journal.pone.0228123

White, A. G., Lees, B., Kao, H.-L., Cipriani, P. G., Munarriz, E., Paaby, A. B., Erickson, K., Guzman, S., Rattanakorn, K., Sontag, E., Geiger, D., Gunsalus, K. C., & Piano, F. (2013). DevStaR: High-throughput quantification of *C. elegans* developmental stages. IEEE Transactions on Medical Imaging, 32(10), 1791–1803. 10.1109/TMI.2013.2265092

Witthames, PR, Thorsen, A, Murua, H, Saborido-Rey, F, Greenwood, LN, Dominguez, R, Korta, M, & Kjesbu, OS. (2009). Advances in methods for determining fecundity: Application of the new methods to some marine fishes. Fishery Bulletin, 107(2), 148–164.

Xu, J., Wang, A., Wang, Y., Li, J., Xu, R., Shi, H., Li, X., Liang, Y., Yang, J., & Gao, T.-M. (2023). AICellCounter: A machine learning-based automated cell counting tool requiring only one image for training. Neuroscience Bulletin, 39(1), 83–88. 10.1007/s12264-022-00895-w

Yang, W., Zhang, X., Tian, Y., Wang, W., & Xue, J.-H. (2019). Deep learning for single image super-resolution: A brief review. IEEE Transactions on Multimedia, 21(12), 3106–3121. 10.1109/TMM.2019.2919431

Yaseen, Z. M., Deo, R. C., Hilal, A., Abd, A. M., Bueno, L. C., Salcedo-Sanz, S., & Nehdi, M. L. (2018). Predicting compressive strength of lightweight foamed concrete using extreme learning machine model. Advances in Engineering Software, 115, 112–125. 10.1016/j.advengsoft.2017.09.004

Zhang, G., Mostad, J. D., & Andersen, E. C. (2021). Natural variation in fecundity is correlated with species-wide levels of divergence in *Caenorhabditis elegans*. G3 Genes|Genomes|Genetics, 11(8), jkab168. 10.1093/g3journal/jkab168

Zhao, Z.-Q., Zheng, P., Xu, S.-T., & Wu, X. (2019). Object detection with deep learning: A review. IEEE Transactions on Neural Networks and Learning Systems, 30(11), 3212– 3232. 10.1109/TNNLS.2018.2876865

