## supplemental materials for "From manual counting to YOLO: Using computer vision to automate large-scale fecundity assays in *C. elegans*"

**Table S1: Hyperparameters evaluated during YOLOv11-L fine-tuning.**

The following hyperparameters were systematically adjusted:

- Optimizer: Algorithm for updating model weights to minimize loss function. Two optimizers were evaluated: Stochastic Gradient Descent (SGD) and Adaptive Moment Estimation with Weight Decay (AdamW).
- IoU (Intersection over Union): Threshold determining the minimum overlap between predicted and ground truth bounding boxes for a prediction to be considered valid.
- HSV_V: Parameter controlling brightness augmentation in the Value channel of the HSV color space, enhancing model robustness to lighting variations.
- Scale: Parameter governing random image resizing during training to improve detection of objects at various sizes.
- Perspective: Parameter controlling viewpoint transformation to enhance generalization across different camera angles.
- Mosaic: Augmentation technique that combines four training images into one, improving detection of small objects and increasing contextual diversity.

*The optimal fine-tuned model utilized the SGD optimizer with the following hyperparameters: IoU=0.4, HSV_V=0.5, scale=0.2, perspective=0.001, and mosaic=0.3.*

| **Hyperparameter** | **Description** | **Values Tested** | **Rationale for Selection** |
|---|---|---|---|
| Optimizer | Algorithm for updating model weights based on loss function gradients | AdamW, SGD | AdamW provides adaptive learning with regularization; SGD offers superior generalization with appropriate momentum parameters |
| IoU | Threshold determining bounding box overlap required for positive detection | 0.3, 0.4, 0.5, 0.6 | Lower thresholds prioritize recall; higher thresholds emphasize precision in detection evaluation metrics |

| Hyperparameter | Description | Values Tested | Rationale for Selection |
| --- | --- | --- | --- |
| HSV_V | Parameter controlling brightness variation in augmentation | 0.1, 0.3, 0.5, 0.7 | Induces illumination invariance, reducing dataset bias toward specific lighting conditions |
| Scale | Factor determining random image resizing during training | 0.2, 0.4, 0.6, 0.8 | Enables development of scale-invariant features through balanced under/over-sampling regimes |
| Perspective | Coefficient controlling viewpoint transformation intensity | 0.0005, 0.001, 0.0015, 0.002 | Simulates viewpoint diversity, enhancing geometric invariance to variation in camera positioning |
| Mosaic | Probability of applying composite image fusion augmentation | 0.3, 0.5, 0.7, 0.9 | Augments contextual diversity and serves as regularization mechanism to improve generalization |

Table S2: Components of the camera imaging station setup.

| **Item** | **Quantity** | **Purchased from** | **Product Number** |
|---|---|---|---|
| Aluminum bottom plate, 12'' x 12'' x 1/2'' | 1 | McMaster-Carr | 8975K135 |
| Acrylic bottom plate, black, 12'' x 12'' x 1/4 '' | 1 | McMaster-Carr | 8505K91/8505K913 |
| Gaffer's tape, black, 1'' wide | 1 | McMaster-Carr | 7612A82 |
| Self-adhesive bumpers, polyurethane, 7/8'' wide, 13/32'' high | 1 | McMaster-Carr | 95495K62, set of 36 |
| Borosilicate glass sheet, 8'' x 8'' x 1/4'' | 1 | McMaster-Carr | 8476K18 |
| 1.5'' diameter post, 14'' long | 1 | ThorLabs | P14 |
| 1.5'' diameter post, 6'' long | 1 | ThorLabs | P6 |
| post clamps | 2 | ThorLabs | C1511 |
| Base plate | 1 | ThorLabs | BA1 |
| 1'' diameter post, 1'' long, 1/4''-20 tap | 1 | ThorLabs | RS1 |
| 1/4''-20 stainless steel hardware kit (5 cap head screws, 2 set screws, 4 washers) | 1 | ThorLabs | HW-Kit2 |
| 3/16'' hex screwdriver | 1 | ThorLabs | |
| Blackout fabric | 1 | Thorlabs | BK5 |
| Devcon Epoxy 5- minute | 1 | Amazon | NSN Stock#8040-00-264-6816: |
| 2-hole inside corner bracket | 1 | Amazon | 80/20 part #4302 |
| 3-12 V Power Supply | 1 | MPJA.com | 36791 PS |
| Red Flexible LED strips, 4.7'' long | 4 | Oznium Inc. | - |
| Camera: 2592 x 1944 pixel sensor, 15 images/s | 1 | Imaging Source | DMK 33GP031 |
| Power Adapter cable for The Imaging Source GigE | 1 | Imaging Source | GigE23/PWR/Trig 1 |
| Power supply for camera | 1 | Imaging Source | PSU 12V/1A/EA 1 Stabilized PSU, EURO and US plug, 100 to 240 VAC, 12VDC, 1A |

| | | | |
|---|---|---|---|
| C-mount extension ring | 1 | Imaging Source | LAex5 |
| C-mount extension ring | 1 | Imaging Source | LAex1 |
| Fujinon lens | 1 | B+H | Fujinon HF12.5SA-1 2/3" 12.5mm f/1.4 C-Mount Fixed Focal Lens |
| Ethernet cable | | Purchased locally | |
| 1/8'' drill bit | | Purchased locally | |
| 5/16'' drill bit | | Purchased locally | |
| Rapid tap heavy duty cutting fluid | 1 | Purchased locally | |

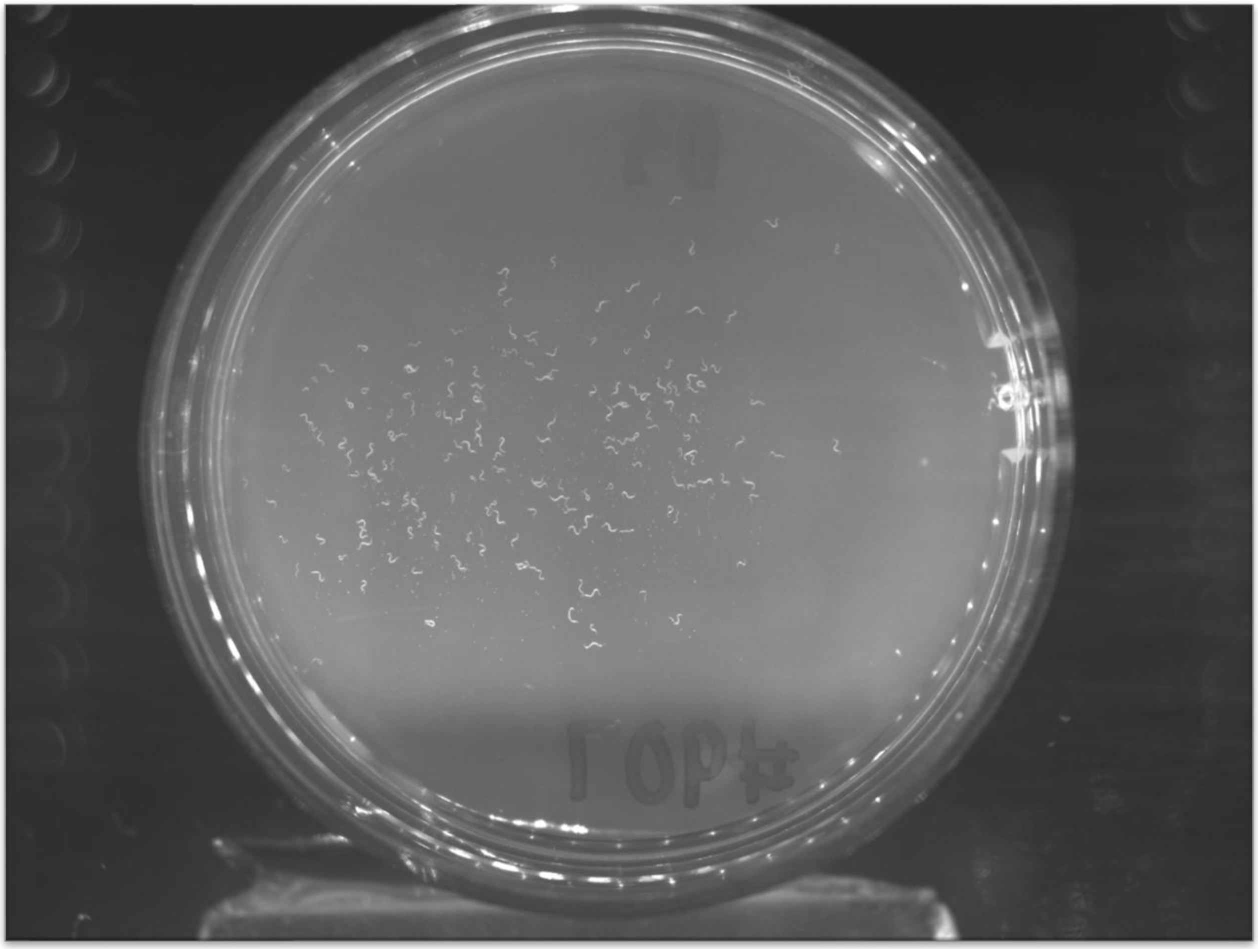

**Figure S1: Example of a raw image in TIFF format.**

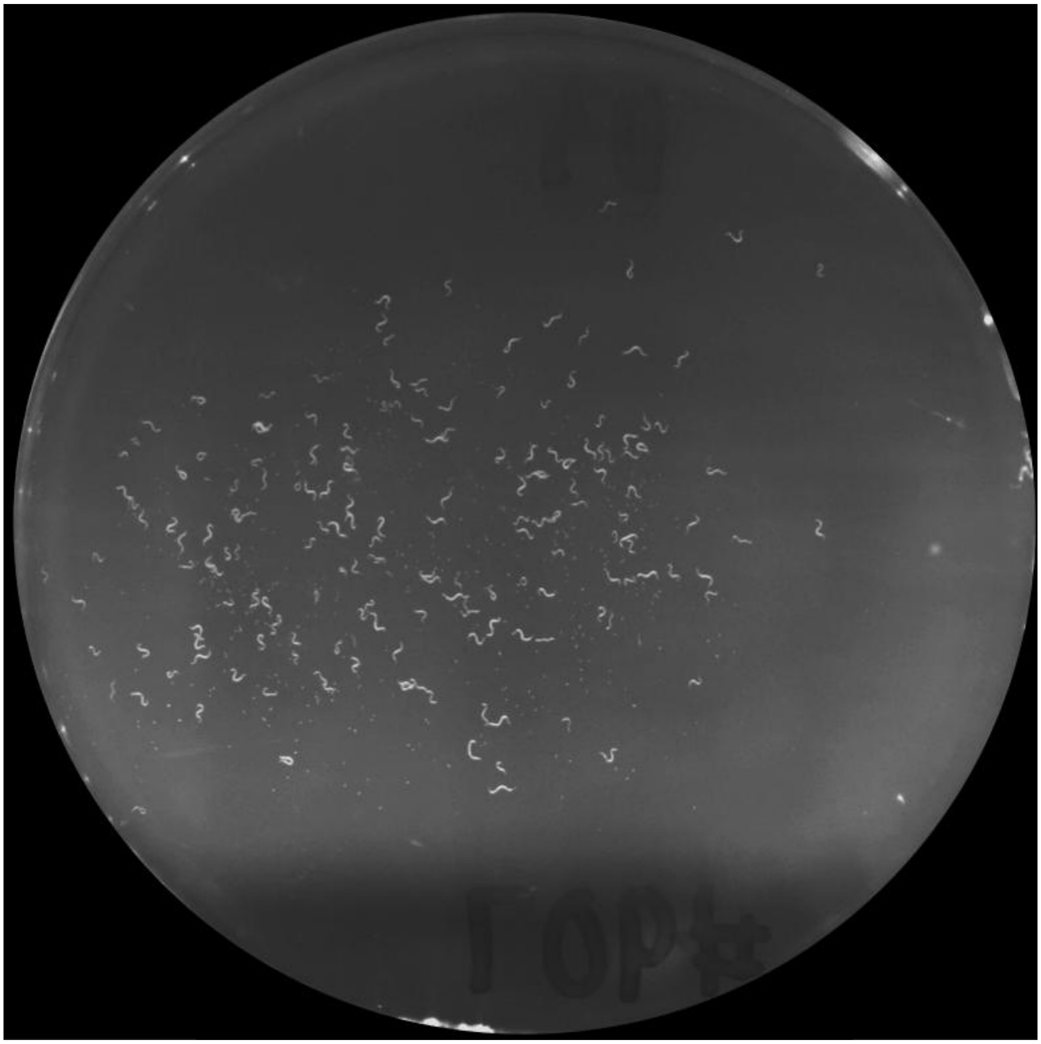

**Figure S2: Example of an augmented image after circular region detection, grayscale conversion and smoothing, and ROI Extraction.**

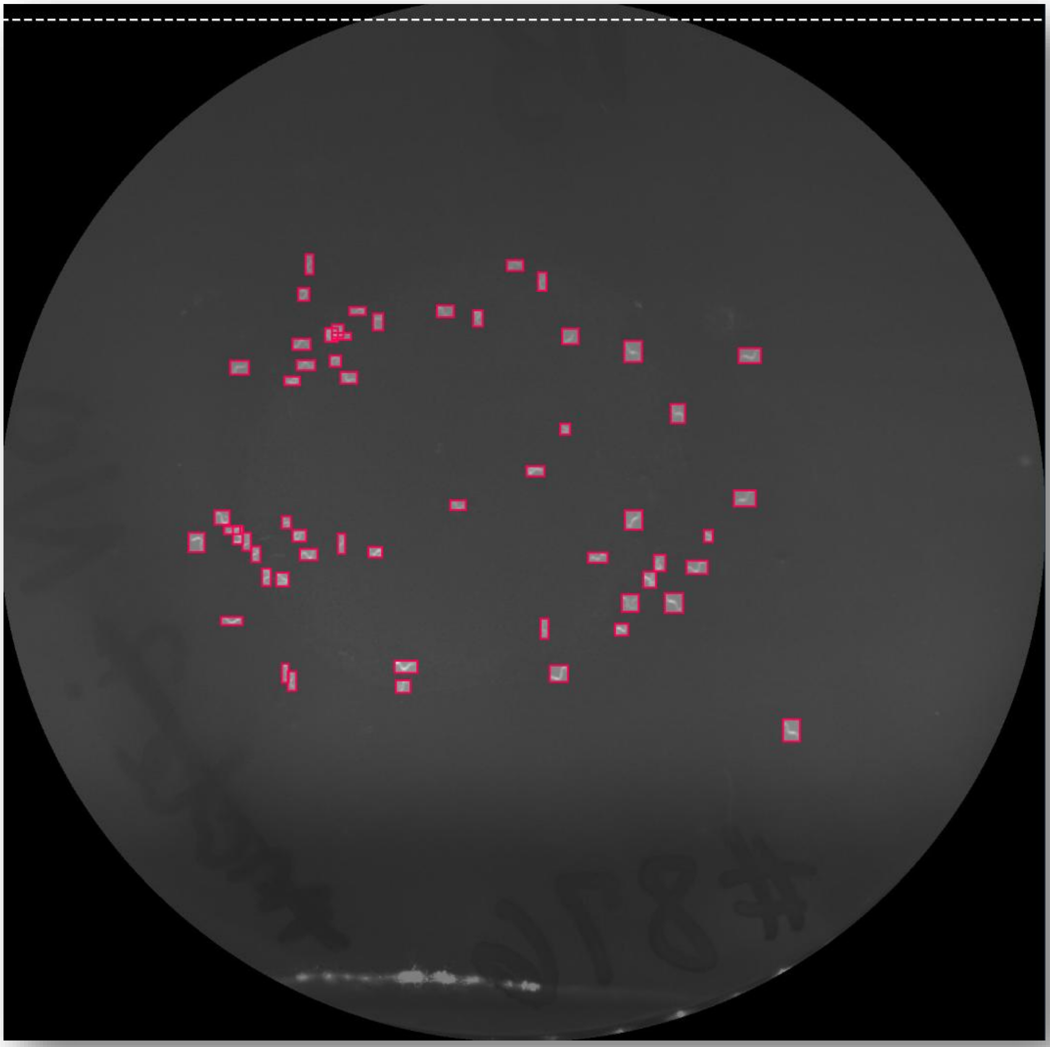

**Figure S3: Example of an image annotated with RoboFlow**

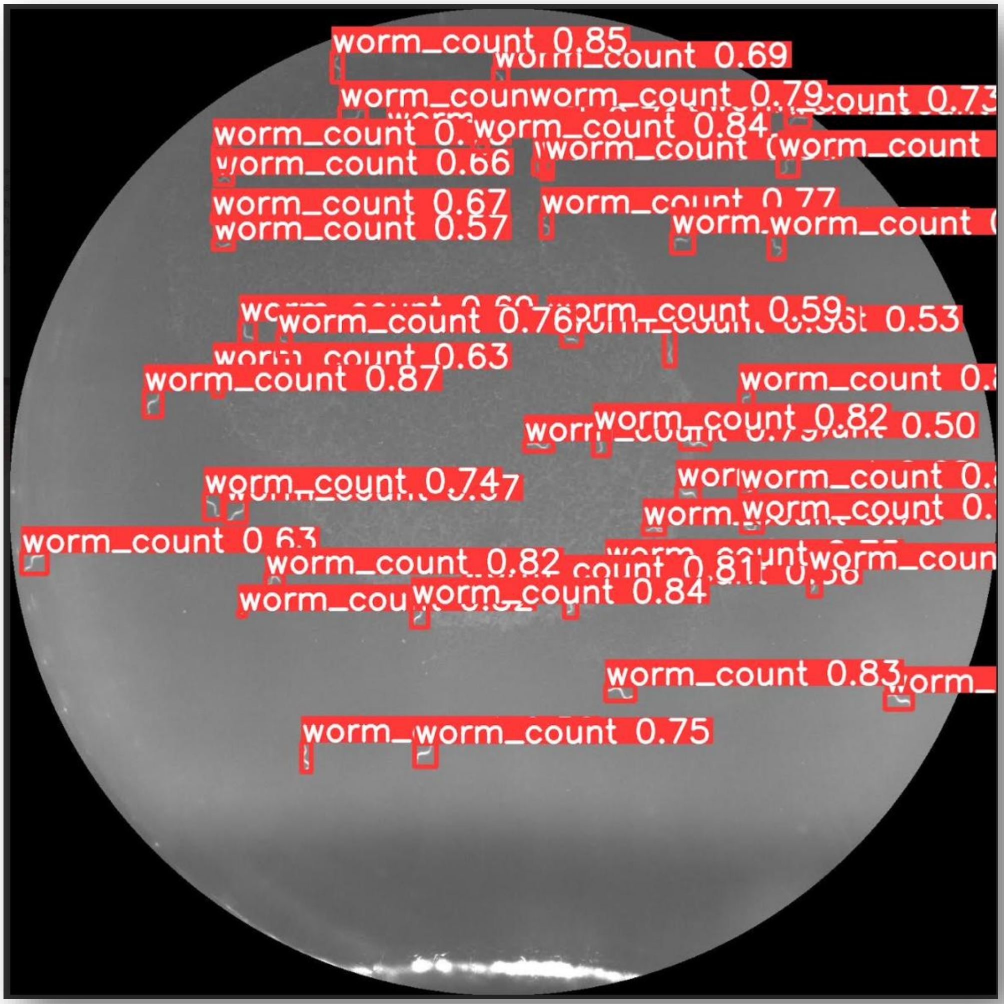


Figure S4: **Example of a predicted Image using the YOLOv11-L-finetuned model with optimal threshold.** All predicted nematodes are highlighted by bounding boxes with their individual score attached.

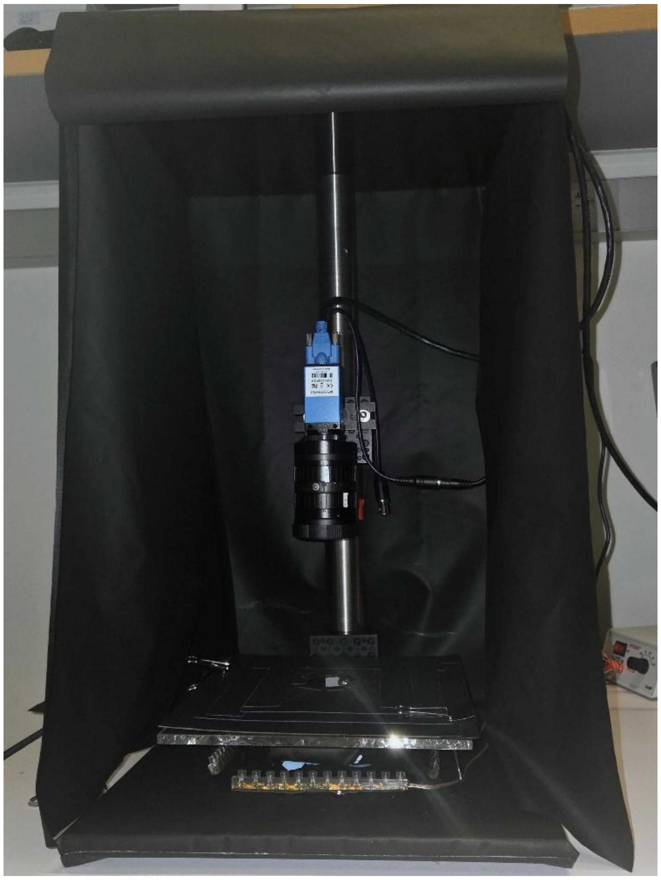

**Figure S5: Imaging station.** The design was adapted from Churgin and Fang-Yen (2015)

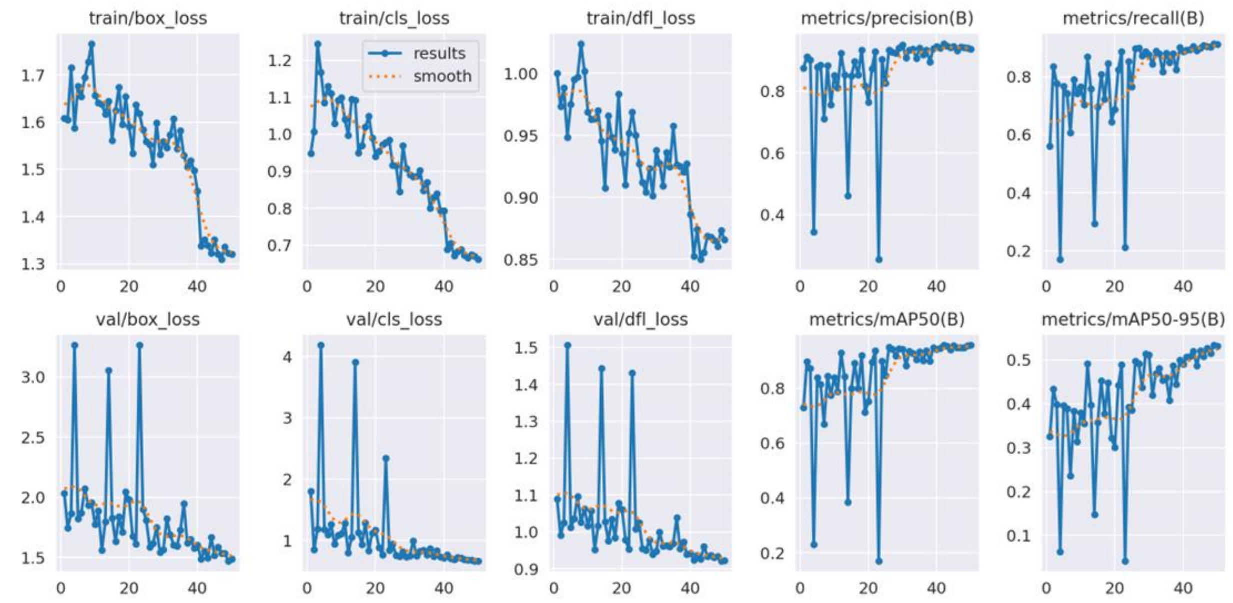


**Figure S6: Example of standard output performance results from the YOLO model (shown for YOLO v11-L).**

Training and validation performance metrics across epochs are shown, including box loss, classification loss, and distribution focal loss (DFL) (top and bottom left panels), as well as precision, recall, mean average precision at IoU $\geqslant$0.5 (mAP50), and mean average precision at IoU $\geqslant$0.5:0.95 (mAP50–95) (right panels). Blue lines indicate raw results, and dashed orange lines indicate smoothed trends.
